# Single-cell Transcriptomics Identifies Gene Expression Networks Driving Differentiation and Tumorigenesis in the Human Fallopian Tube

**DOI:** 10.1101/2020.05.28.119933

**Authors:** Huy Q. Dinh, Xianzhi Lin, Forough Abbasi, Robbin Nameki, Marcela Haro, Claire E. Olingy, Heidi Chang, Lourdes Hernandez, Simon A. Gayther, Kelly N. Wright, Paul-Joseph Aspuria, Beth Y. Karlan, Rosario I. Corona, Andrew Li, BJ Rimel, Matthew Siedhoff, Fabiola Medeiros, Kate Lawrenson

## Abstract

The human fallopian tube harbors the cell-of-origin for the majority of high-grade serous ‘ovarian’ cancers (HGSCs), but its cellular composition, particularly the epithelial component, is poorly characterized. We performed single-cell transcriptomic profiling in 12 primary fallopian specimens from 8 patients, analyzing around 53,000 individual cells to map the major immune, fibroblastic and epithelial cell types present in this organ. We identified 10 epithelial sub-populations, characterized by diverse transcriptional programs including SOX17 (enriched in secretory epithelial cells), TTF3 and RFX3 (enriched in ciliated cells) and NR2F2 (enriched in early, partially differentiated secretory cells). Based on transcriptional signatures, we reconstructed a trajectory whereby secretory cells differentiate into ciliated cells *via* a RUNX3^high^ intermediate. Computational deconvolution of the cellular composition of advanced HGSCs based on epithelial subset signatures identified the ‘early secretory’ population as a likely precursor state for the majority of HGSCs. The signature of this rare population of cells comprised both epithelial (*EPCAM, KRT*) and mesenchymal (*THY1*, *ACTA2*) features, and was enriched in mesenchymal-type HGSCs (P = 6.7 × 10^−27^), a group known to have particularly poor prognoses. This cellular and molecular compendium of the human fallopian tube in cancer-free women is expected to advance our understanding of the earliest stages of fallopian epithelial neoplasia.

## Introduction

The fallopian tube serves as a conduit between the ovary and the uterus to allow the transport of the fertilized egg for implantation in the endometrium. It is derived from the Müllerian ducts, which also give rise to the uterus, cervix, and upper vagina (Healey, 2012). The fallopian tube is a dynamic organ that undergoes structural modifications in response to the menstrual cycle, menopause and tubal ligation (Crow et al., 1994; Donnez et al., 1985). The fallopian tube is composed of three layers: the mucosa, a seromuscular layer, and the serosa. The mucosa consists of the lamina propria and the epithelium. Within the lamina propria resides a network of fibroblast-mesenchymal cells (Wang et al., 2015a) and immune cells. T cells represent the dominant population of lymphoid cells found in the fallopian tube, with CD8+ T cells being the prevalent cell type (Ardighieri et al., 2014). Neutrophils are the second most prevalent population of immune cells, and are relatively more numerous in the fallopian tube compared to the rest of the female reproductive tract (Givan et al., 1997; Smith et al., 2006).

Three histologic cell types comprise the epithelium: ciliated, secretory, and intercalary cells. Ciliated cells are responsible for oocyte pickup, embryo transport, and tubal fluid flow. The intercalary cells are thought to be a population of epithelial stem-like cells, which may play a role in the initiation of serous tumors (Paik et al., 2012). The secretory cells comprise approximately 60% of the epithelial cells found in the fallopian tube, and secrete growth factors, nutrients, antimicrobial agents and anti-cellular stress agents into the tubal fluid. Fallopian tube secretory epithelial cells are of particular interest as there is now evidence that the majority of high-grade serous ‘ovarian’ cancers (HGSCs) originate from the secretory cells of the distal fallopian tube, with the recent development of a step-wise progression model starting from a p53 signature which progresses into serous tubal intraepithelial carcinoma (STIC), and finally to HGSC (Lee et al., 2007; Reade et al., 2014; Wang et al., 2015b). Secretory cell expansion and loss of ciliated cells are thought to be early events in the development of serous cancers (Wang et al., 2015b), but molecular drivers of these phenomena are unknown, as the differentiation path for tubal epithelia is poorly understood. Here we employed droplet-based single cell RNA-sequencing (scRNA-seq) to map the cellular composition of the human fallopian tube. We identify epithelial subpopulations, including early and transitioning populations, and the critical transcriptional networks that define cellular populations in this organ. We then leverage the signatures of epithelial subsets to identify putative cells-of-origin for HGSCs.

## Results

### Single cell transcriptomic analyses of normal human fallopian tubes

We performed single cell RNA-sequencing (scRNA-seq) of twelve human fallopian tube tissues from eight cancer-free individuals (Figure 1A). The phenotypic and clinical characteristics are summarized in Table 1. Three specimens were from the distal, fimbriated region where the tube is in closest proximity to the ovary; the remaining nine specimens were from mid-portion of the organ. In one patient we profiled matched ampulla, infundibulum and fimbrial portions of the fallopian tube. Patient age ranged from 31-62 years; all patients except patient 8 were premenopausal. The interval between onset of last menstrual period to surgery ranged from 3-31 days. All patients in the cohort were undergoing gynecological surgeries for benign conditions including leiomyoma (7 patients), adenomyosis (3 patients) and endometriosis (2 patients). Two patients were parous (each with two live births). None of the patients were known or suspected to be carriers of any genetic abnormalities associated with hereditary cancer predisposition syndromes. In total we profiled 64,042 single cells isolated from the 12 specimens. We then excluded cells with ≤ 200 genes expressed and/or high mitochondrial RNA content (mtRNA ≥ 10%). Potential doublets were removed on a sample-wise basis based on the number of captured cells. After this filtering, a total of 52,908 cells remained for analysis. The average number of genes surveyed was 1,083 per cell (range: 709-1,636), and the average number of reads per cell was 96,166 (range: 31,578-259,280).

**Figure 1.**
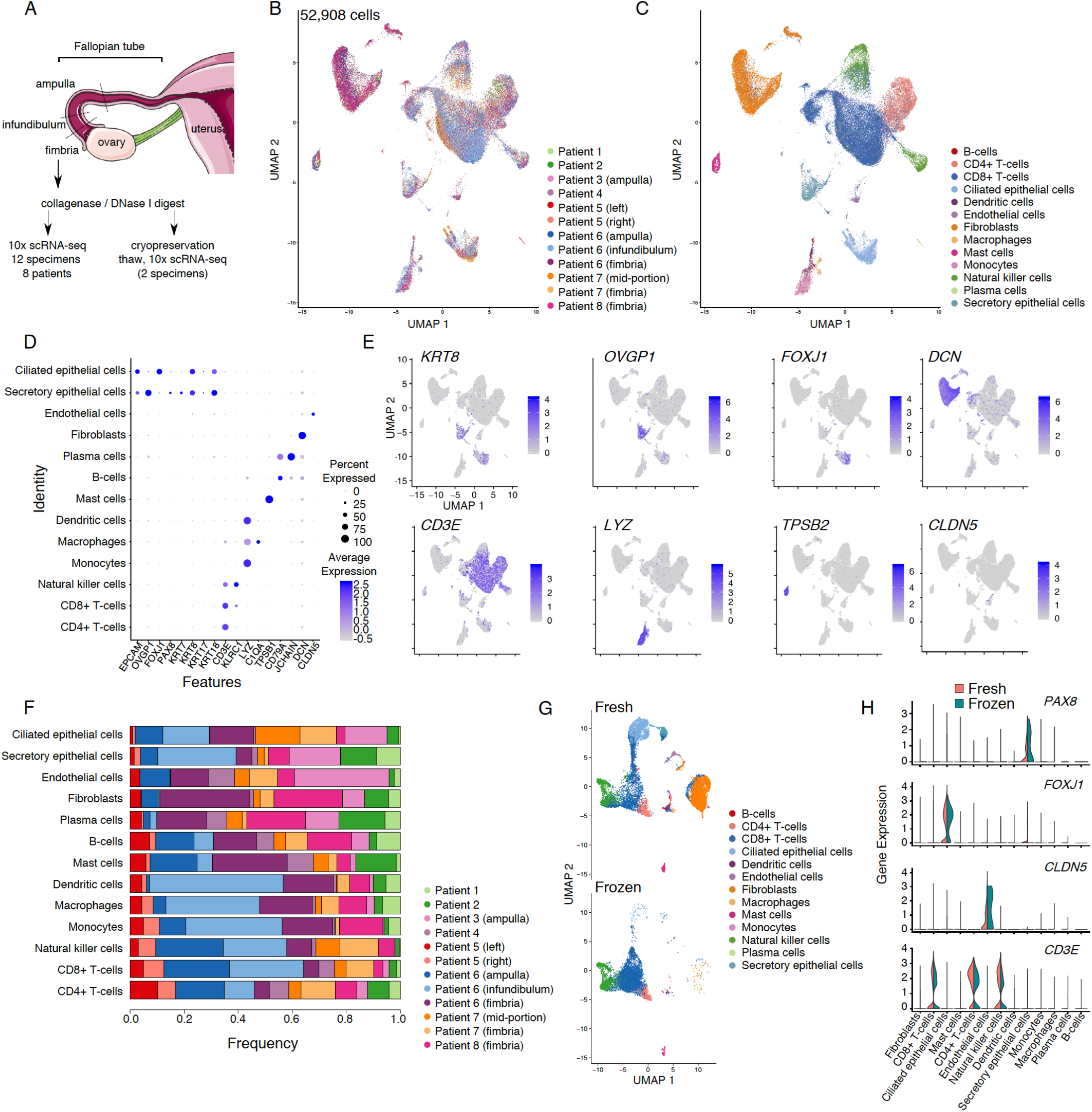
A cellular gene expression atlas of human fallopian tubes in cancer free women. (A) Schematic showing overall study design. (B) UMAP plot showing the major cellular clusters by patient and (C) by cell type. (D) Scaled expression level of cell-type defining markers and percentage of positive cells in each cluster. (E) UMAP plots showing cluster-specific expression patterns of representative cell-type defining markers. (F) Frequency of cells in each cluster, by patient. (G) UMAP plots for 2 specimens before and after cryopreservation. (H) Violin plot of marker expression in 2 specimens, before and after cryopreservation. UMAP, Uniform Manifold Approximation and Projection.

These data were integrated and batch corrected using canonical correlation-based alignment analysis implemented in the Seurat package (Stuart et al., 2019). Using Louvain-based clustering method, we detected 13 major cellular populations based on well-defined markers (Methods) (Figure 1D-E). In the immune microenvironment of the tube, the major populations were lymphoid T cells including CD4 and CD8 T cells and natural killer cells. Myeloid cells (monocytes, macrophages, dendritic cells and mast cells), B cells and plasma cells were detected at lower frequencies. We also detected a large population of fibroblasts/smooth muscle cells and a small population of endothelial cells. Epithelial cells formed two distinct clusters, one comprised of *PAX8/OVGP1*-expressing secretory epithelial cells, and the other consisting of *FOXJ1*-positive ciliated epithelia (Figure 1C-E). All major clusters were detected in each of the specimens analyzed (albeit at different frequencies) (Figure 1F). To determine the impact of cryopreservation on fallopian tube tissues, cells from two specimens from patients 6 and 7 were viably frozen in 10% DMSO/90% fetal bovine serum, then thawed around 6 months later and subjected to a second round of capture and sequencing. Gene signatures from the 13 major cell types present in the fresh tissue profiles were used as a reference to predict the cell type labels in the two frozen samples. Even though lower cell numbers were captured in these specimens compared to the fresh primary tissues, we were able to recover all the major cell types after cryopreservation (Figure 1G, Figure S1). All the major immune cellular populations were present in the repeat sample, T cells were the most conserved, and myeloid cells (monocytes, macrophages, and dendritic cells) were depleted from cryopreserved samples. Epithelial cells and fibroblasts, although present, were also depleted following cryopreservation (Figure S1). Expression of key cell-type defining markers was stable following thawing (Figure 1H). Taken together, these results indicate that freeze-thaw cycles do not significantly impact the expression profile of fallopian tube cells, and that lymphocytes within dissociated fallopian tube tissues are particularly resilient to cryopreservation.

We asked if scRNA-Seq could reveal transcription factors (TFs) associated with major cell types in the fallopian tube by implementing the SCENIC pipeline (Aibar et al., 2017). SCENIC integrates co-expression network analysis and motif-based analyses to identify TFs and downstream genes (called regulons) that characterize the distinct transcriptional programs that define the major cell clusters (Figure 2A, Figure S2). Transcriptional signatures of fibroblasts were enriched for multiple regulons including KLF4, EGR1, GATA2 and TCF4. Epithelial factors included SOX17, BARX2 and BHLHE41 in secretory epithelial cells, plus KLF5, TTF3 and RFX3 in ciliated epithelial cells. We recently identified SOX17 as a master regulator in high-grade serous ovarian cancers (Reddy et al., 2019), but SOX17 is not known to be a marker in physiologically normal fallopian tube epithelial cells. To validate this finding we performed immunohistochemistry for SOX17 on 14 fallopian tubes from cancer-free women; strong SOX17 staining was observed in all specimens, and was independent of age, menopausal status, and *BRCA1/2* mutation status (Figure 2B, Table S1). As expected, SOX17 staining tended to have greater intensity in secretory cells, compared to ciliated cells.

**Figure 2.**
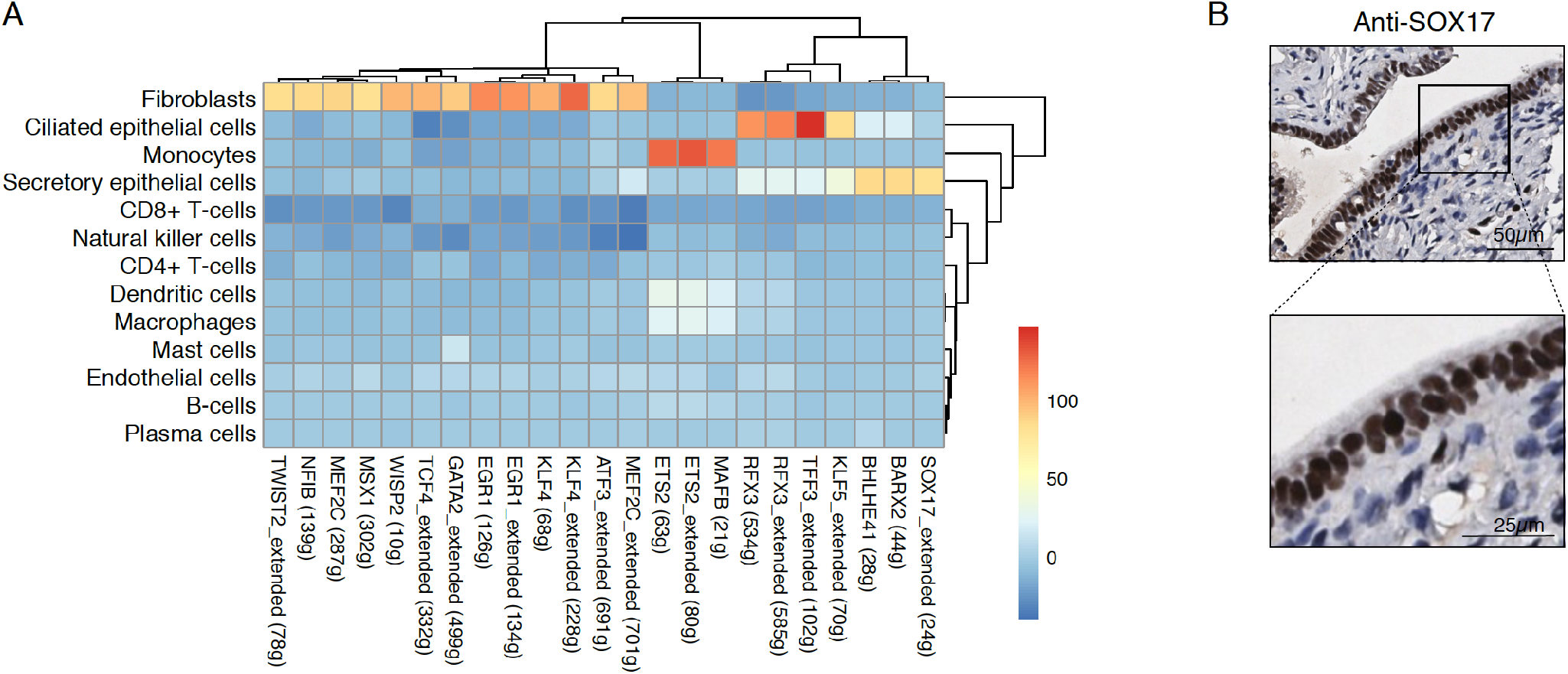
Transcription factor analyses in human FT cell populations. (A) Identification of TF regulons enriched in specific cell types using SCENIC (t-value from a linear model test of regulon activity score of one subset *versus* the rest, see Methods). Numbers in brackets denote the number of genes contained in the regulon, “extended” means the regulons include motifs that have been linked to the TF using motif similarity from SCENIC pipeline. (B) Validation of SOX17 expression in human FT tissues using immunohistochemistry.

### Epithelial components of the fallopian tube

As fallopian tube epithelial cells, particularly the secretory epithelial cells, are the likely precursors of most HGSCs (Karst et al., 2011; Lee et al., 2007; Perets et al., 2013; Reade et al., 2014; Wang et al., 2015b), we focused on defining the epithelial landscape of the fallopian tube. Epithelial cell populations were identified by expression of well-defined markers *EPCAM*, keratin genes (*KRT7, KRT8* and *KRT18*), *OVGP1* and *FOXJ1*; cells expressing at least one of these markers were annotated as epithelial. Clustering was performed on 4,592 epithelial cells (8.7% of the total cell population), using *PAX8/OVGP1* and *FOXJ1* as defining markers of secretory and ciliated epithelial cells, respectively. Integration analysis stringently required each epithelial subset to represent at least 2% of cells in at least 3 samples. Using these criteria, we identified a total of 10 epithelial cell clusters, which we categorized as *PAX8^high^* and *OVGP1^high^* secretory epithelial cells (3 clusters, secretory 1-3); *FOXJ1^high^* ciliated epithelial cells (4 clusters, ciliated 1-4); and three clusters that lowly expressed *PAX8*, *OVGP1* and *FOXJ1* and could not therefore be classified as secretory or ciliated (unclassified 1-3) (Figure 3A). Across the 10 epithelial subpopulations, there was extensive heterogeneity in the expression of known FT epithelial markers. *EPCAM* was highly expressed in secretory cluster 2 and ciliated clusters 1, 2 and 4, but was low/absent in the remaining clusters. Secretory clusters 2 and 3 and ciliated cluster 2 expressed high levels of low molecular weight keratins 7, 8 and 18 (Figure 3B). Unclassified cluster 3, secretory clusters 2 and 3, and ciliated 4 expressed *KRT10*; *KRT17* was expressed in unclassified 3, secretory cluster 2 and ciliated cluster 2. Overall, ciliated epithelial cells were more abundant than secretory cells (47.3%, range 11.4-83.6% for ciliated epithelial cells, *versus* 38.4% range 3.4-84.2% for secretory epithelial cells), although this difference was not significant (P = 0.31, two-tailed paired t-test) (Figure 3C). Secretory clusters 2 and 3 and ciliated clusters 1, 2 and 4 each comprised ~10% of the epithelial cells captured. Cells from all samples contributed to each cluster (Figure S3).

**Figure 3.**
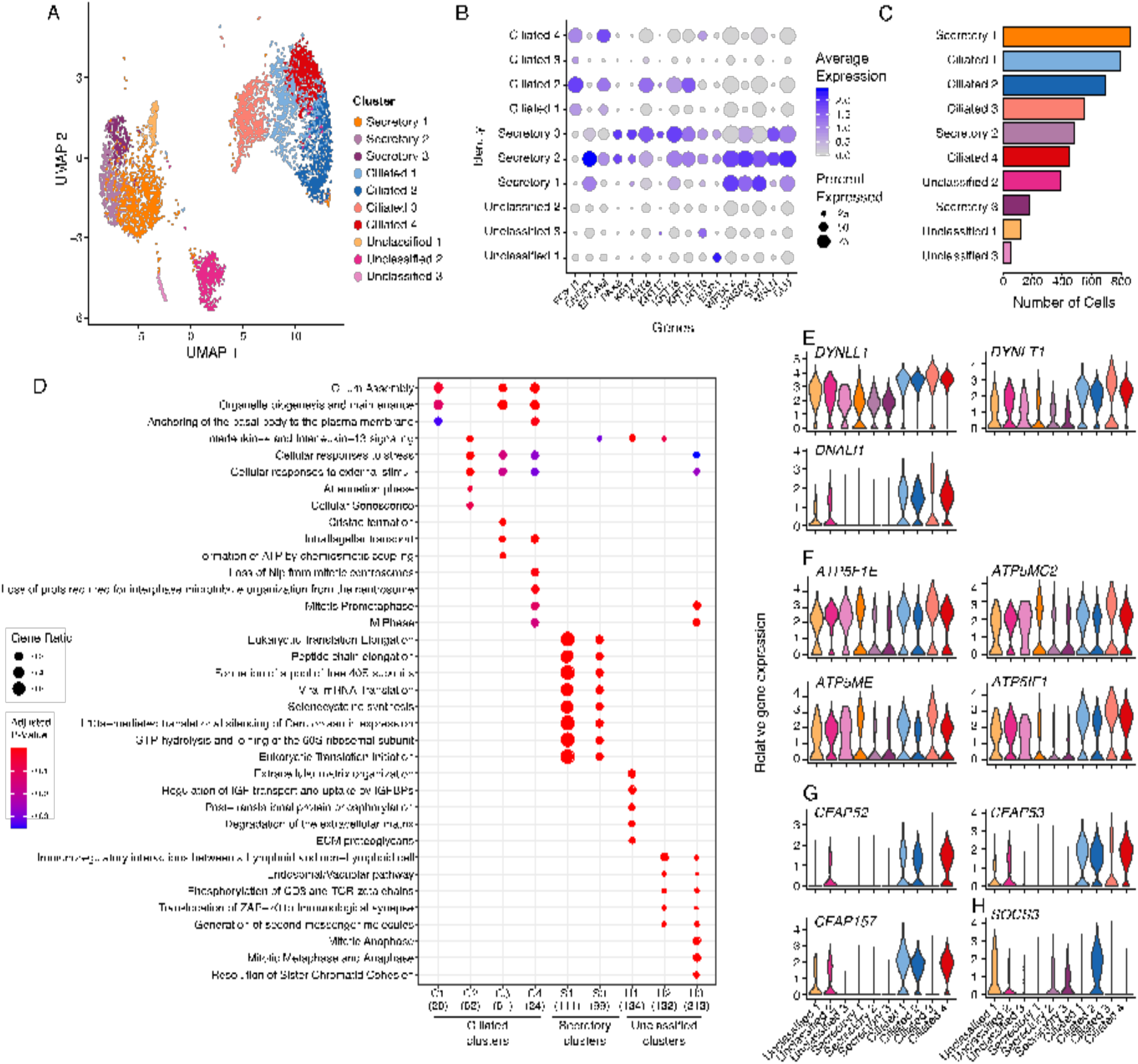
Epithelial cell subpopulations in the human fallopian tube. (A) UMAP representation of 10 distinct epithelial clusters revealed by graph-based Louvain clustering implemented in the Seurat single cell data analysis package. (B) Dot plot presentation of scaled expression of epithelial, secretory and ciliated cell markers. (C) Frequency histogram for the 10 epithelial clusters. (D) Pathway analyses performed using the ReactomePA R package. All pathways with P < 0.05 are shown. Secretory cluster 2 has zero enriched pathways reaching P < 0.05. Expression of (E) dyenin protein-encoding genes, (F) ATP synthase 5 subunit genes and (G) genes encoding cilia proteins. UMAP, Uniform Manifold Approximation and Projection.

We performed differential gene expression analyses within each epithelial cell subtype to identify differentially expressed genes across the subpopulations of fallopian epithelial cells (average log fold change (logFC) > 0.5, FDR adjusted p-value < 0.01, and 25% of cells within the cluster expressing the gene of interest) (Table S2). Fallopian tube secretory epithelial cell (FTSEC) clusters were defined by higher expression of *PAX8* and *OVGP1* compared to ciliated epithelial cells (Figure 3B). When we surveyed the expression of known biomarkers, secretory clusters 2 and 3 exhibited the highest expression of *PAX8*, *KRT7* and *ESR1*; and clusters 1 and 2 expressed relatively higher levels of *OVGP1* and *WFDC2* (which encodes ovarian cancer biomarker HE4) compared to cluster 3. FTSECs expressed high levels of less well known markers including cysteine rich secretory protein 3 (*CRISP3*) - a protein that has been implicated in epithelial cell proliferation and adhesion in the endometrium (Evans et al., 2015) but not in the fallopian tube; secretory leukocyte protease inhibitor (*SLPI*), expressed by HGSCs (Cancer Genome Atlas Research Network, 2011); mesothelin (*MSLN*), a proposed target for immunotherapy in ovarian cancer (Haas et al., 2019); and secreted chaperone clusterin (*CLU*) (Figure 3B). Secretory cluster 1 was the most abundantly represented epithelial population in our data set, with, on average, 23.2土19.0% of the epithelial cells in each specimen assigned to this cluster (range 1.1-56.5%). This cluster was defined by 129 genes that were strongly enriched in myriad pathways associated with translation elongation (adjusted P-value = 3.8 × 10^−155^, Table S3), reflecting the high secretory activity of these cells (Figure 3D). Similar pathways were enriched in the genes that defined secretory cluster 3 (on average 2.8% of cells per sample), albeit with more modest p-values. This cluster expressed *KRT17*, which was absent in clusters 1 and 2 and likely corresponds to the *KRT17*-positive secretory cluster (C4) recently described in tubal epithelium from women with gynecologic cancers (Hu et al., 2020).

Ciliated clusters 1, 2 and 4 consisted of differentiated ciliated cells (Figure 3D, Table S3). Cluster 3 highly expressed genes encoding dynein components (*DYNLT1*, *DNALI1* and *DYNLL1*) (Figure 3E), and genes associated with energy production, with six subunits of mitochondrial ATP synthase 5 overexpressed (Figure 3F). Cluster 3 expressed some genes encoding cilia and flagella associated proteins (*CFAP53* and *CFAP126*) but other genes encoding cilia and flagella associated proteins were absent/lowly expressed (*CFAP157*, *CFAP73*, *CFAP52* and *CFAP126*) (Figure 3G), suggesting the cells in this cluster are not yet fully differentiated into a ciliated phenotype. In contrast, ciliated cluster 4 represented differentiated ciliated epithelial cells and expressed high levels of genes encoding dyenin components in addition to genes encoding cilia-associated proteins (Figure 3G). Pathways preferentially enriched in ciliated cluster 2 included cellular responses to stress (adjusted P-value = 7.02 × 10^−5^), senescence (adjusted P-value = 0.008) and p53 regulated cell death (adjusted P-value = 0.04) (Figure 3E, Table S3). In this cluster suppressor of cytokine signaling 3 (*SOCS3*) was specifically expressed at high levels; this is a cytokine inducible gene that negatively regulates cytokine signaling (Figure 3H) that may be involved in senescence of terminally differentiated ciliated cells (Mallette et al., 2010).

### Reconstructing differentiation trajectories in the human fallopian tube

Pseudotime analysis was performed using Monocle2, to order epithelial cells based on expression patterns (Qiu et al., 2017). This analysis constructed a cellular trajectory whereby secretory cells differentiate into ciliated cells *via* unclassified clusters 2 and 3 (Figure 4A, Figure S4A). Unclassified cluster 1 and secretory cluster 3 were enriched at the earliest time points and may represent putative progenitor cells and/or committed cells early in the differentiation process, before progressing into secretory cluster 2 and then 1. Unclassified cluster 1 exhibited low expression of all of the genes that define ciliated and secretory epithelial cells (Figure 3B) and expressed *CD44* and *EPCAM* (Figure 4B), known features of tissue stem cells in many tissue types, including the fallopian tube (Paik et al., 2012). This cluster was defined by a set of 186 genes that were enriched in pathways associated with the extracellular matrix and responses to growth factors (Figure 3E). We searched for cell surface markers unique to this population and identified *THY1*, which encodes CD90 (Figure 4B). We validated the EPCAM^hi^/CD44^hi^/THY1^hi^ population in an independent fallopian tissue specimen using flow cytometry. Single cells were isolated using the sample protocol implemented for single cell RNA-seq. CD45+ cells were depleted before performing flow cytometry with fluorescently-tagged antibodies targeting EPCAM, CD44 or THY1. A triple positive population comprised 4.7% of the EPCAM-positive cells in this specimen (Figure 4C). In the scRNA-seq this cluster also expressed high levels of *IGFBP5*, *LGALS1* (which encodes galectin-1) and *ACTA2* (which encodes smooth muscle actin) (Figure 4B) and was the cluster with the highest expression of *ESR1* (encoding the estrogen receptor) (Figure 3B).

**Figure 4.**
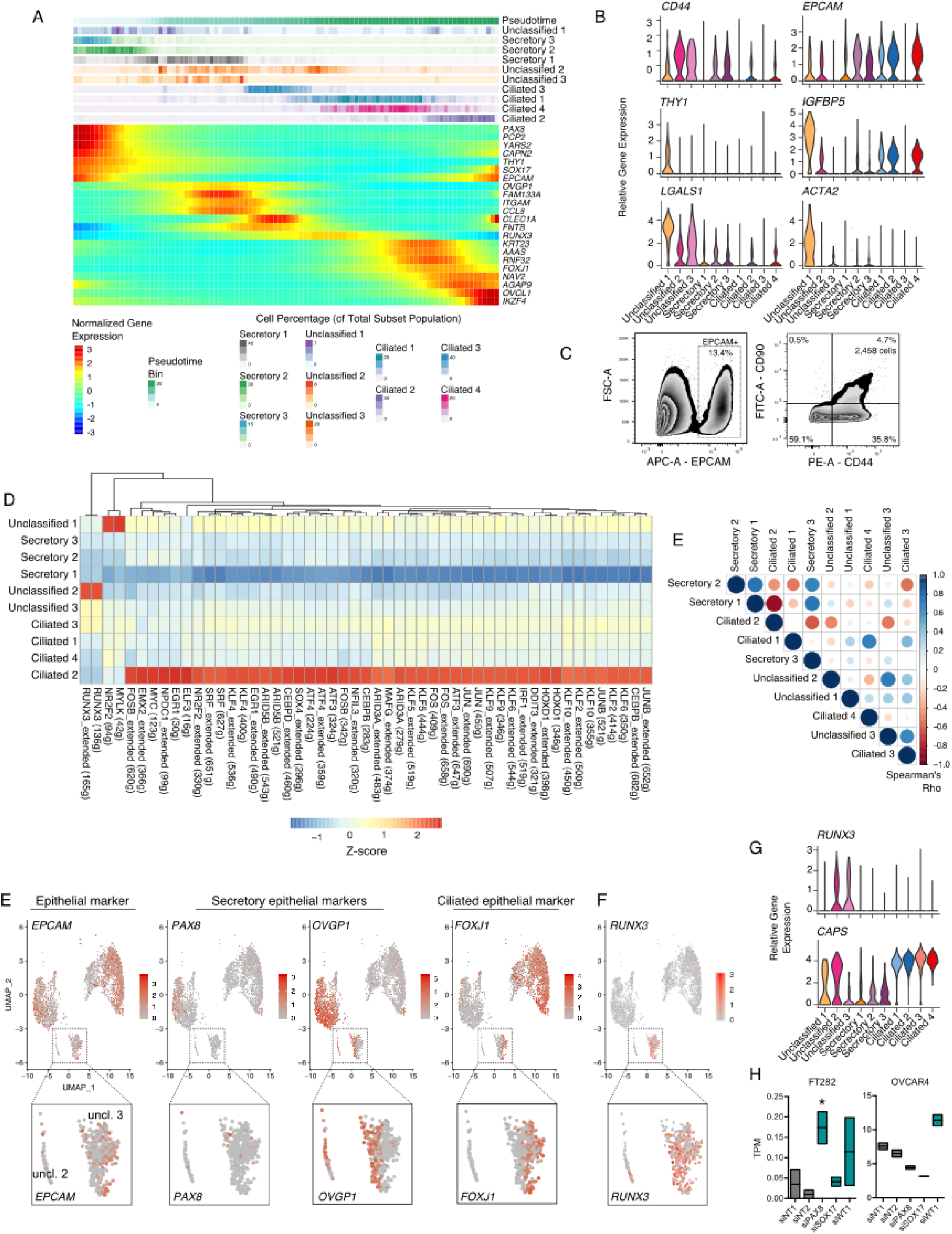
A differentiation trajectory for fallopian tube epithelia. (A) Pseudotime analyses performed using Discriminative Dimensionality Reduction with Trees (DDRTree) implemented in the Monocle2 package. The upper portion of the panel shows inferred pseudotime, divided into 100 bins, from early (left) to late (right). Proportion of cells from each cluster assigned to each bin are shown below. The lower panel shows expression of known and novel biomarkers across the pseudotime trajectory identified using a differential expression test (linear model of pseudotime) in Monocle2. (B) Expression of *CD44*, *EPCAM*, and markers enriched in unclassified cluster 1. (C) Validation of the presence of unclassified cluster 1 in a fallopian tube tissue using flow cytometry. (D) Transcription factor regulon analyses, performed using SCENIC. (D) Correlation of t-values from linear model test contrasting 1 subset vs the other 9 subsets. Color of dot denotes direction of correlation (Spearman correlation test), size denotes correlation coefficient. (E) UMAP plots showing expression of *EPCAM*, and secretory/ciliated cell genes. Inset panels show a gradient of *PAX8*, *OVGP1* and *FOXJ1* expression in unclassified clusters 2 and 3. (F) *RUNX3* expression is restricted to unclassified clusters 2 and 3. (G) Violin plots of *RUNX3* and *CAPS* expression. (H) Floating box plots showing *RUNX3* expression after *PAX8*, *SOX17* or *WT1* knockdown. Upper and lower boundaries of the box denote maximum and minimum expression values, horizontal line denotes mean. TPM, transcripts per million, normalized RNA-seq reads. N = 2 independent siRNA transfections. * |log FC| >1, P = 0.02, negative binomial regression performed using DEseq2. UMAP, Uniform Manifold Approximation and Projection.

Transcriptional regulation network analyses, again performed using the SCENIC pipeline, highlighted the NR2F2, EMX2 and MYC regulons as enriched in the early secretory unclassified 1 subset. Each of these factors has been previously implicated in stem/progenitor cell biology (Figure 4D) (Cheng et al., 2010; Kozak et al., 2020; Takahashi and Yamanaka, 2006). These regulons were highly specific to this early, partially differentiated state. We observed a strong inverse correlation between TF regulons enriched in differentiated ciliated cells (ciliated cluster 2) and differentiated secretory cells (secretory cluster 1) (Spearman’s correlation coefficient = −0.87) (Figure 4E) and across the secretory and ciliated subsets, a pattern of TF regulon activity that mirrored the pseudotime trajectory. Unclassified cluster 2, which was located at the interface of secretory and ciliated cell phenotypes, showed high activity of the RUNX3 cistrome (Figure 4D). The RUNX3 regulon was also activated, albeit to a lesser extent, in unclassified cluster 3 and ciliated cluster 3. Unclassified cluster 2 expressed a gradient of decreasing *OVGP1* and increasing *FOXJ1* along the pseudotime trajectory and likely represents a population of cells undergoing the transition from a secretory to a ciliated phenotype (Figure 4F). On average 8.0% of epithelial cells were assigned to this cluster (standard deviation 5.3%; range: 0.0-15.4%). Unclassified cluster 3 comprised rare cells representing, on average, 1.8% of all epithelial cells in each specimen (s.d. = 1.5, range = 0.2 - 5.2%). This cluster was characterized by decreasing expression of *OVGP1* and mitotic pathways and genes were strongly enriched (Figure 3E, Figure S4B), even though we regressed out cell cycle phase scores during pre-processing to minimize the impact of cell cycle heterogeneity on cluster calls. This cluster likely corresponds to cluster C9 from Hu *et al.* (2020). Cells in this cluster appeared to represent an early transitioning cell population at the point of losing *OVGP1* expression. Unclassified cluster 2 has begun to acquire some features of ciliated cells, including *CAPS* expression (Figure 4G) and then the pathway for ciliated differentiation progresses from ciliated 3 (which expresses dynein genes but few cilia genes) into ciliated clusters 1 and 4 - two clusters of fully differentiated ciliated cells. Ciliated cluster 2, which comprises differentiated ciliated cells undergoing cell death and senescence, represents the endpoint in the differentiation trajectory (Figure 3D, Figure 4A, Table S3).

Consistent with the RUNX3 regulon being specifically enriched in transitioning unclassified clusters 2 and 3, *RUNX3* gene expression was unique to these clusters (Figure 4D,F). TFs that characterize fallopian tube secretory epithelial cells are PAX8 (Levanon et al., 2010; Ozcan et al., 2011), SOX17 (this study and Reddy et al., 2019) and WT1, a clinically used biomarker of HGSC (Köbel et al., 2008; Shimizu et al., 2000). To understand the relationship of RUNX3 to these major secretory cell transcription factors, we transiently knocked down *PAX8*, *SOX17* and *WT1* in normal fallopian tube epithelial cells (FT282) and a HGSOC cell line (OVCAR4). *RUNX3* expression was almost undetectable in the FT282 cells, but following depletion of *PAX8*, *RUNX3* expression increased 3.0-fold (P = 0.02, negative binomial regression performed using DEseq2) (Figure 4H, Figure S4C-E). *RUNX3* expression was not significantly altered following depletion of *SOX17* or *WT1* (log_2_ FC = 0.78, P = 0.67; FC = 2.3, P = 0.14). Knockdown of *PAX8* and *SOX17* in the OVCAR4 cell line modestly down-regulated *RUNX3* expression (log_2_ FC = −0.66, p = 8.50×10^−7^; log_2_ FC = −0.73, P = 1.29 × 10^−7^), whereas *WT1* depletion lead to a moderate increase in *RUNX3* expression (log_2_ FC = 0.46, P = 1.6 × 10^−4^) (Figure 4H). We note that the expression changes observed in the OVCAR4 tumor model did not reach the threshold of absolute log_2_ FC > 1 typically applied to identified differentially expressed genes in RNA-seq data. This suggests that in the normal physiological state, PAX8 inhibits *RUNX3* expression, but that this regulation is attenuated during tumorigenesis.

### Early secretory cell signatures are enriched in ex vivo fallopian tube cultures and in high-grade serous cancers

We derived signatures of the 10 epithelial clusters, combining secretory clusters 2 and 3, and ciliated clusters 1, 3 and 4 due to overlap in the signature genes (Figure 5A). Informed by the characteristics described above, we re-annotated ‘Unclassified cluster 1’ as ‘early secretory’, while unclassified clusters 2 and 3 were denoted as ‘transitioning’ clusters. Using the MuSiC method (Wang et al., 2019) with our combined signatures (see Methods), 6 epithelial signatures were used to deconvolute bulk RNA-seq profiles of 68 FTSEC primary cultures (Lawrenson et al., 2019), to infer the cell subpopulations that preferentially grow *in vitro*. In 25 out of 68 cultures (39.4%), the ‘early secretory’ signature was strongly enriched (at least top 25% percentile of estimated frequency, i.e ≥ 0.7 frequency). A subset of 21 samples with higher expression of *OVGP1* were most enriched for the differentiated secretory cell signature (top 25% percentile, i.e. frequency > 0.34; Figure 5B). Signatures of secretory cluster 1 and ciliated cells were poorly enriched overall. To see if this pattern is maintained after cellular immortalization with ectopic expression of *TERT* and mutant *TP53,* we applied cell type prediction *via* the label transfer function implemented in Seurat, to map cellular identities from all 10 epithelial subsets in scRNA-seq profiles for the FT282 cell line (Karst et al., 2014; Lawrenson et al., 2019). Consistent with the results seen for the *ex vivo* cultures, in this immortalized cell line the majority of cells were labeled as ‘early secretory’ (705 out of 1,690 total cells, 41.7%) (Figure S5). These data imply that early secretory cells preferentially grow *in vitro*, or that *in vitro* culture results in a partial de-differentiation of secretory epithelial cells that is maintained following immortalization.

**Figure 5.**
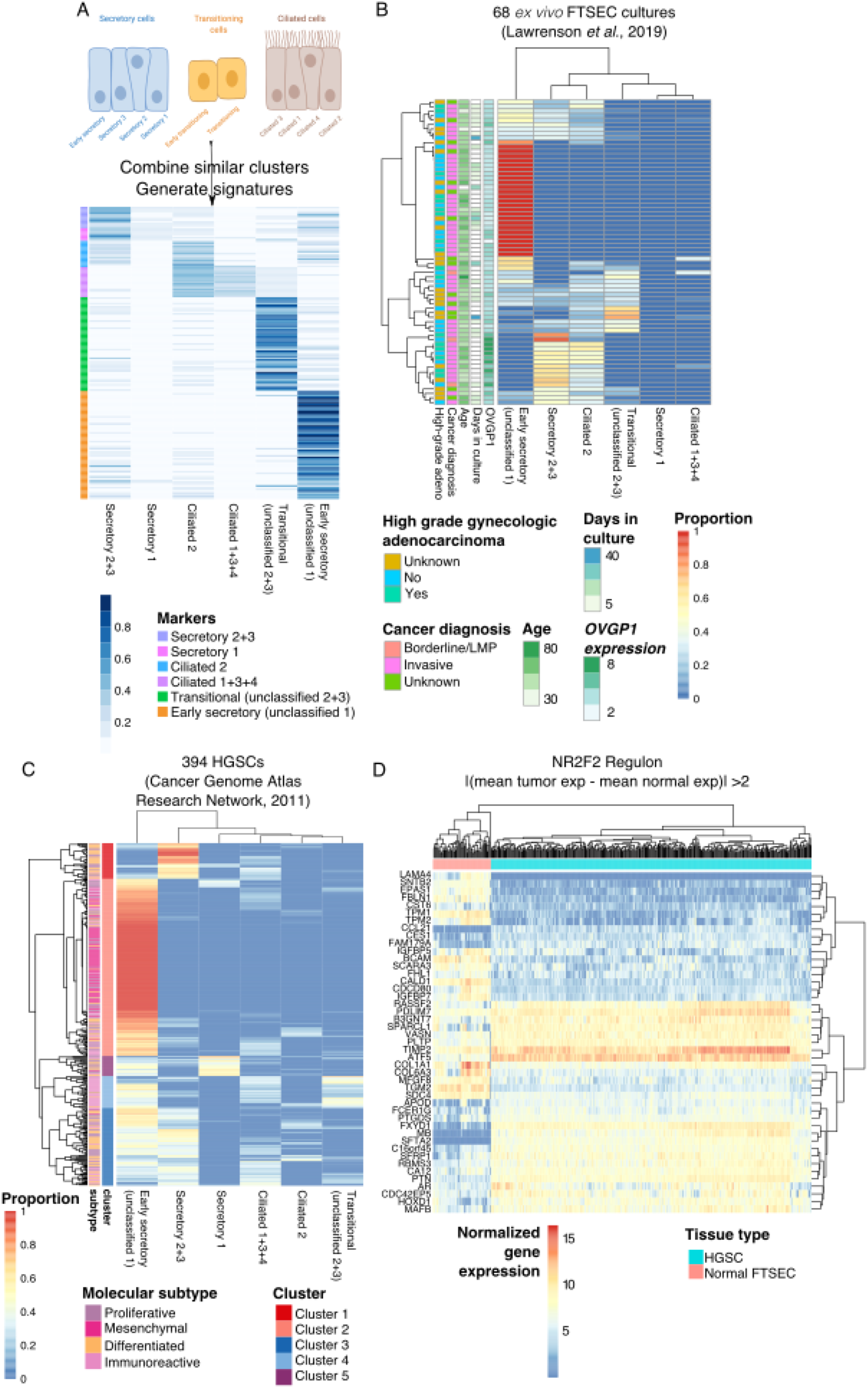
Signatures of early secretory epithelia are enriched in fallopian epithelial cultures and HGSCs. (A) Development of epithelial subset signatures informed by pseudotime analyses; subsets with similar profiles of marker genes were combined for computational deconvolution analysis. (B) Deconvolution of bulk RNA-seq profiles of 68 *ex vivo* FTSEC cultures: heatmap of the composition of 6 subsets in each sample. A subset of patients had diagnoses of high-grade adenocarcinomas of the ovary, endometrium or cervix. (C) Deconvolution of 394 HGSCs from TCGA. (D) The NR2F2 regulon is dysregulated in HGSCs compared to FTSECs. Top differentially expressed genes were identified by calculating |(mean HGSC expression - mean FTSEC expression)| >2. On all heatmaps rows and columns are ordering using unsupervised hierarchical clustering based on Euclidian distances.

The six signatures were then applied to 394 HGSC RNA-seq profiles generated by The Cancer Genome Atlas. Unsupervised clustering of tumors based on the deconvoluted cell-type composition identified five clusters of tumors with similar composition (Figure 5C). As with the normal FTSEC cultures, the ‘early secretory’ signature was the most strongly enriched in the majority of tumors (cluster 2, 204 out of 394 cases), and was particularly enriched in tumors of the mesenchymal molecular subgroup (89/92 mesenchymal samples were in cluster 2; P = 6.7 × 10^−27^, hypergeometric test) (Table S4) (Cancer Genome Atlas Research Network, 2011; Tothill et al., 2008). Clusters 1 and 3 were characterized by tumors exhibiting the differentiated secretory 2+3 signature, respectively; both were enriched with the differentiated molecular subgroup (P = 1.4 × 10^−7^ and P = 1.2 × 10^−5^, respectively, hypergeometric test). Tumors in cluster 4 were enriched for the transitional cell signature and the immunoreactive TCGA subtype was overrepresented in this cluster (P = 1.6 × 10^−18^, hypergeometric test) Cluster 5 was comprised of tumors with signatures of secretory cluster 1, and was enriched in the proliferative molecular subtype (P = 0.01). In sum, the gene signatures of epithelial subsets were enriched within each TCGA molecular subtypes and the computational deconvolution highlights the association of those gene signatures with established molecular subtypes.

Finally, since the NR2F2 regulon was activated in the early secretory cluster (Figure 5D), we tested whether this regulon was dysregulated during tumorigenesis using our data set of 68 normal FTSEC *ex vivo* cultures integrated with 394 HGSCs from TCGA (Lawrenson et al., 2019). Ninety-three genes from the NR2F2 regulon were expressed in our data set (Figure S6). Unsupervised clustering based on expression of these genes clearly separated tumor cells from normal cells, with dysregulation of genes involved in negative regulation of cell proliferation (q-value = 0.001) such as *AR*, *EPAS1* and *SPARCL1* and regulation of cell-substrate adhesion, including *LAMA4*, *CCL21* and *BCAM* (Figure 5D, Figure S6).

## Discussion

Single cell gene expression analyses bridge the interface of cell biology, pathology and gene regulation. These analyses are transforming our understanding of intra-tissue heterogeneity in development, normal tissue homeostasis, and disease alike (Cao et al., 2019; Wang et al., 2020). Using single cell RNA-seq performed with the 10x Genomics platform, we generated a cellular atlas of the human fallopian tube in cancer-free women, and identified the transcriptional networks driving the expression programs of 13 major cell types. While the fallopian tube is located internally, the genital tract is exposed to the external environment, and thus a complex network of lymphocytes, macrophages and dendritic cells to control invading pathogens. The immune landscape of this organ was dominated by T cells, which were largely composed of CD8+ T cells, consistent with previous measurements by immunohistochemistry (IHC) (Ardighieri et al., 2014). Reproductive tract T cells have variable cytolytic capacity throughout the menstrual cycle, suggesting their function is hormonally regulated (White et al., 1997). The phenotype and function of fallopian tube T cells, particularly during tumorigenesis, will be of great interest for designing effective immunotherapies. While previous work using IHC found CD68+ macrophages to be as abundant as CD8+ T cells in 10 non-cancerous fallopian tube samples, we measured a very low frequency (1.4% of leukocytes) of macrophages using scRNA-Seq. This difference could be attributed to challenges in extracting macrophages from the extracellular matrix during sample preparation (Givan et al., 1997). Prior work using flow cytometry also identified neutrophils as one of the predominant leukocyte subtypes in healthy fallopian tubes (Givan et al., 1997; Smith et al., 2006), but IHC was unable to detect substantial numbers of extravascular neutrophils (Ardighieri et al., 2014). Similarly, in our analysis, we did not detect neutrophils, although they are generally challenging to capture in scRNA-Seq data, probably due to their tendency to rapidly undergo apoptosis *ex vivo* and low RNA yield.

We identified 10 epithelial subpopulations present in the fallopian tube and projected a cellular trajectory based on the transcriptional signatures of each cluster (Figure 6). The model for differentiation pathway for tubal epithelia was challenged in recent years with the demonstration that, rather than secretory and ciliated cells deriving from a bipotent progenitor, in murine models all FT epithelia derives from a unipotent PAX8+ precursor, with secretory cells giving rise to ciliated cells (Ghosh et al., 2017). Our data provide evidence that this model holds true in human tissues. We identified two clusters that were enriched early in the epithelial cellular trajectory and appear to correspond to partially differentiated, late progenitor populations (Snegovskikh et al., 2014). The first was *KRT17*-positive secretory cluster 3, and the second, the “early secretory” cluster (former unclassified cluster 1), which cluster was characterized by EPCAM^hi^, CD44^hi^ and THY1^hi^ expression. The early secretory cluster expressed both epithelial (*EPCAM*, *KRT19*) and mesenchymal (*THY1*, *DCN*) characteristics as well as smooth muscle actin and estrogen receptor, and thus bore some similarities to myoepithelial cells of the uterus. This population expresses *IGFBP5* and *LGALS1*, genes implicated in homeostasis of stem and/or progenitor populations in normal and diseased tissues (Byrd et al., 2019; Garcés et al., 2020; Tumbar et al., 2004; You et al., 2018) and *MEG3*, an imprinted long noncoding RNA. Previous reports have identified an enrichment of stem cells in the fimbrial portion of the tube (Paik et al., 2012). In our one patient with matched ampulla, infundibulum and fimbrial tissue, the proportion of THY1+ cluster cells was highest in the fimbrial portion of the tissue (10% *versus* <3% in the mid and proximal regions). The specimen with the highest proportion of THY1 early secretory cluster came from an individual with a diagnosis of endometriosis. Endometriosis was not observed in the portion of the tissue that was subjected to pathology review, although we cannot rule out the presence of microscopic foci of endometriosis in the segment of tissue subjected to scRNA-seq. Nonetheless it is plausible that the altered pelvic microenvironment associated with endometriosis may impact epithelial differentiation in the fallopian tube. Larger studies will be required to test these hypotheses.

**Figure 6.**
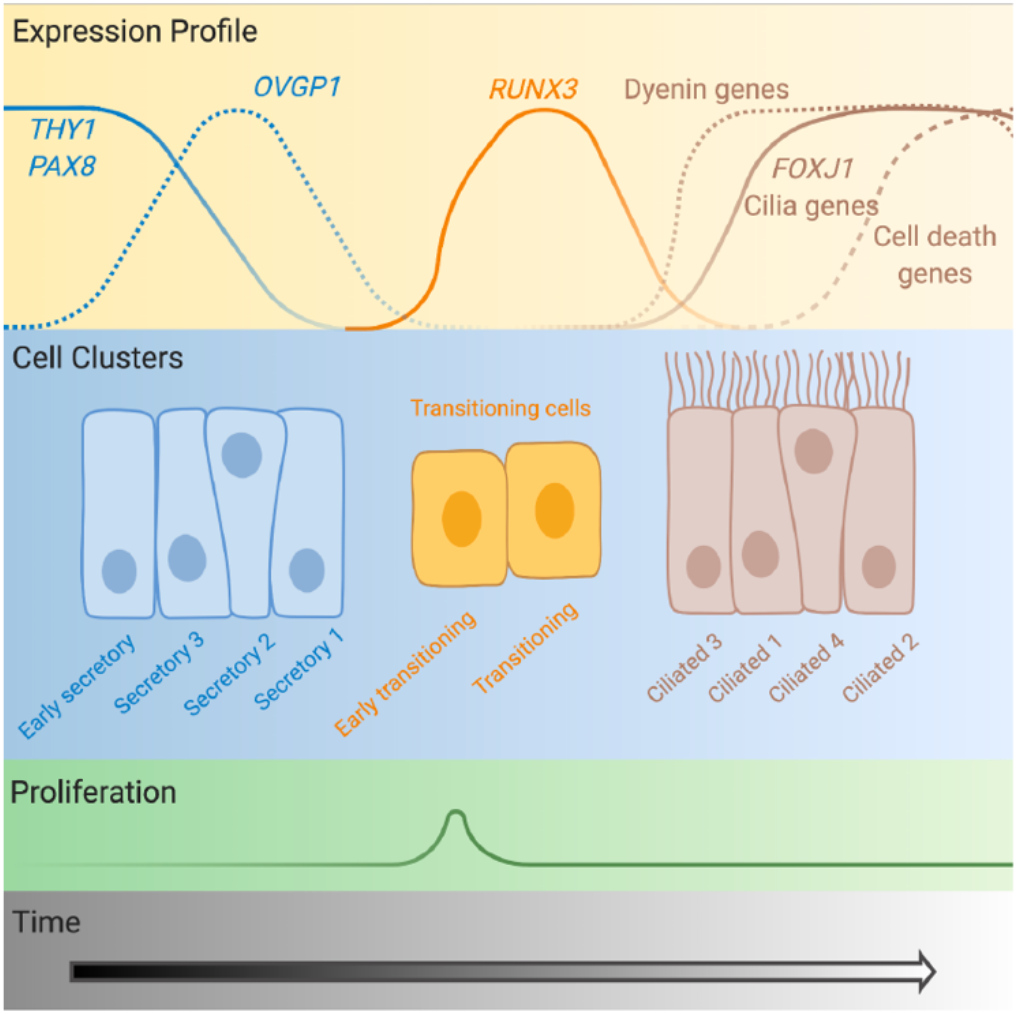
Diagram of the cellular differentiation path for fallopian tube epithelia. Expression profiles of key markers and transcription factors are shown (not to scale). The ‘early secretory’ cluster corresponds to the former ‘unclassified cluster 1’, the ‘early transitioning’ cluster corresponds to ‘unclassified cluster 3’, the ‘transitioning’ cluster corresponds to ‘unclassified cluster 2’. Created with BioRender.com.

Epithelial cell dynamics in the fallopian tube have not been extensively studied, but in the last two decades, interest in this organ has been piqued by strong evidence to support a fallopian tube origin for the majority of high-grade serous ‘ovarian’ carcinomas (Callahan et al., 2007; Chen et al., 2017; Lee et al., 2007; Medeiros et al., 2006; Meserve et al., 2017). The pseudostratified epithelium of the fallopian tube consists of two major cellular populations. A secretory population that expresses the PAX8 transcription factor is thought to be the predominant cell-of-origin for HGSC (Callahan et al., 2007; Chen et al., 2017; Ducie et al., 2017; Lawrenson et al., 2019; Lee et al., 2007; Medeiros et al., 2006; Meserve et al., 2017). Fallopian secretory epithelia are more proliferative *in vitro* and *in vivo* and less competent at repairing DNA damage compared to the FOXJ1-positive ciliated population (Levanon et al., 2010). Our results suggest that an early secretory cell subset preferentially adapts to culture and is characterized by a molecular signature that is maintained in advanced tumors. Whether this specific cellular population represents a precursor of HGSC, or whether a more differentiated cell dedifferentiates during neoplastic transformation has yet to be determined. Since the presence of long stretches of secretory cells, secretory cell outgrowths (SCOUTs) is thought to be an early step in the development of HGSC (Meserve et al., 2017), it is likely that a perturbation in the differentiation path for tubal epithelia is an early event in high-grade serous ‘ovarian’ carcinogenesis. We identified RUNX3 as a putative regulator of a transitional state representing an intermediate between secretory and ciliated phenotypes. RUNX3 reportedly promotes proliferation and chemoresistance of ovarian cancer cells *in vitro* (Barghout et al., 2015; Lee et al., 2011; Nevadunsky et al., 2009). In our analyses RUNX3 was overexpressed in tumor cells compared to fallopian tube cells, but repression by Müllerian TFs was context-specific, suggesting regulation of RUNX3 is perturbed during tumorigenesis. Since an increased ratio of secretory to ciliated cells represents a more cancer-prone state, this suggests that loss of the normal transition of secretory to ciliated phenotype is an early step in neoplastic transformation; more work is needed to elucidate whether RUNX3 plays a key role in the development of SCOUTs and STICs, as our data suggest.

One of the greatest insights emerging from this work is the identification of transcription factors regulating the different stages of tubal epithelial cell differentiation. SOX17 was enriched in secretory cells, this factor had not been implicated in HGSC or fallopian biology until we recently described this protein as a master transcription factor for HGSC and both a regulator of and binding partner for PAX8 (Reddy et al., 2019; Moreira and Drapkin, personal communication). NR2F2 and EGR1 regulons were enriched in ‘early secretory’ and ‘secretory 1’ clusters. We also observed a striking inverse correlation in the regulons enriched in differentiated endpoint for ciliated cells (ciliated cluster 2) and the most differentiated secretory cell cluster (secretory 1). Across the other clusters, activation state of these regulons mirrored the cellular trajectory inferred by pseudotime analyses. This suggests that TF networks required for differentiation of ciliated cells are largely suppressed in the differentiated secretory cell state. Master regulators underlying this negative feedback likely include *PAX8*, *RUNX3* and *FOXJ1*, or interactions with these factors. Perturbation of this fine-tuned transcriptional regulation is likely an early step in tumor development.

We compared our epithelial subsets with the recently published work using SMART-Seq2 of EPCAM-positive fallopian tube cells (Hu et al., 2020). We were able to match all of the subsets they identified using the gene signatures from their work. Our secretory cluster 3 had features of both the KRT17+ and inflammatory clusters, our unclassified cluster 2 likely corresponds to their ‘cell cycle’ cluster, and our ciliated cluster expressed genes that corresponded to the ‘stress’ gene set defined in this prior study. Our early secretory cluster expressed both genes from the differentiated subset (*DPP4*/*LTBP4*/*SLC25A25*) and EMT subset (*ACTA2*/*TIMP3*), although we note that the Hu *et al.,* study did not attempt to describe putative progenitor cells in the tube. The overall consistency between studies indicates that the epithelial subpopulations described are stable in women of different ages and diagnoses, independent of the technology used. Larger studies will be required to identify significant perturbations in epithelial subpopulation frequencies and transcriptional signatures associated with age and lifetime ovulatory cycles, both major risk factors for HSGC. Our study has some limitations. While it is known that fallopian tube fimbriae likely harbor the cell of origin for the majority of HGSCs, the small sample size of our study precluded in-depth analyses of features specific to the fimbrial portion of the organ, to gain insight into why these cells are more prone to neoplastic transformation. In addition, analyses of cells collected from women at high risk of ovarian cancer, such as women with germline mutations in the *BRCA/2* genes, are expected to be the most informative for discovery of biomarkers early stages of pelvic serous carcinoma, but were not included in the current study as the extensive pathological examination of tubes prophylactically removed from *BRCA1/2* mutation carriers (Medeiros et al., 2006) currently limits the application of scRNA-seq to these specimens. Finally, we were underpowered to explore the impact of the fluctuating steroid hormones due to phases in the menstrual cycle, or menopause.

While the data presented here will be valuable for the study of infertility and benign conditions such as pelvic inflammatory disorder and ectopic pregnancy, the greatest translational impact is likely to come from new insights into origins and early development of HGSC. We observed that the fallopian epithelial cells that grow *ex vivo* and immortalized cultures represent an early differentiated fallopian tube secretory epithelial cell type and are therefore highly relevant for cancer studies, as the signature of these THY1+ early secretory cells was enriched in a majority of HGSCs. This signature was the most enriched in mesenchymal-type tumors, perhaps unsurprisingly, given the mesenchymal properties of this population, and suggests that further studies are warranted to characterize this population in the high-risk setting. The majority of patients with HGSC are diagnosed with advanced stage disease and have an overall 5-year survival rate of less than 40%, and there is a pressing need to study normal tubal tissues to understand the earliest stages of disease to develop more effective strategies for early detection and prevention. We expect this atlas of cells, transcriptomes and transcription factor networks in disease-free tissues will lead to translational insights by laying the foundation for identifying subtle alterations in the composition and functioning of the fallopian tube in the context of disease or elevated cancer risk.

## Methods

### Key Resources Table

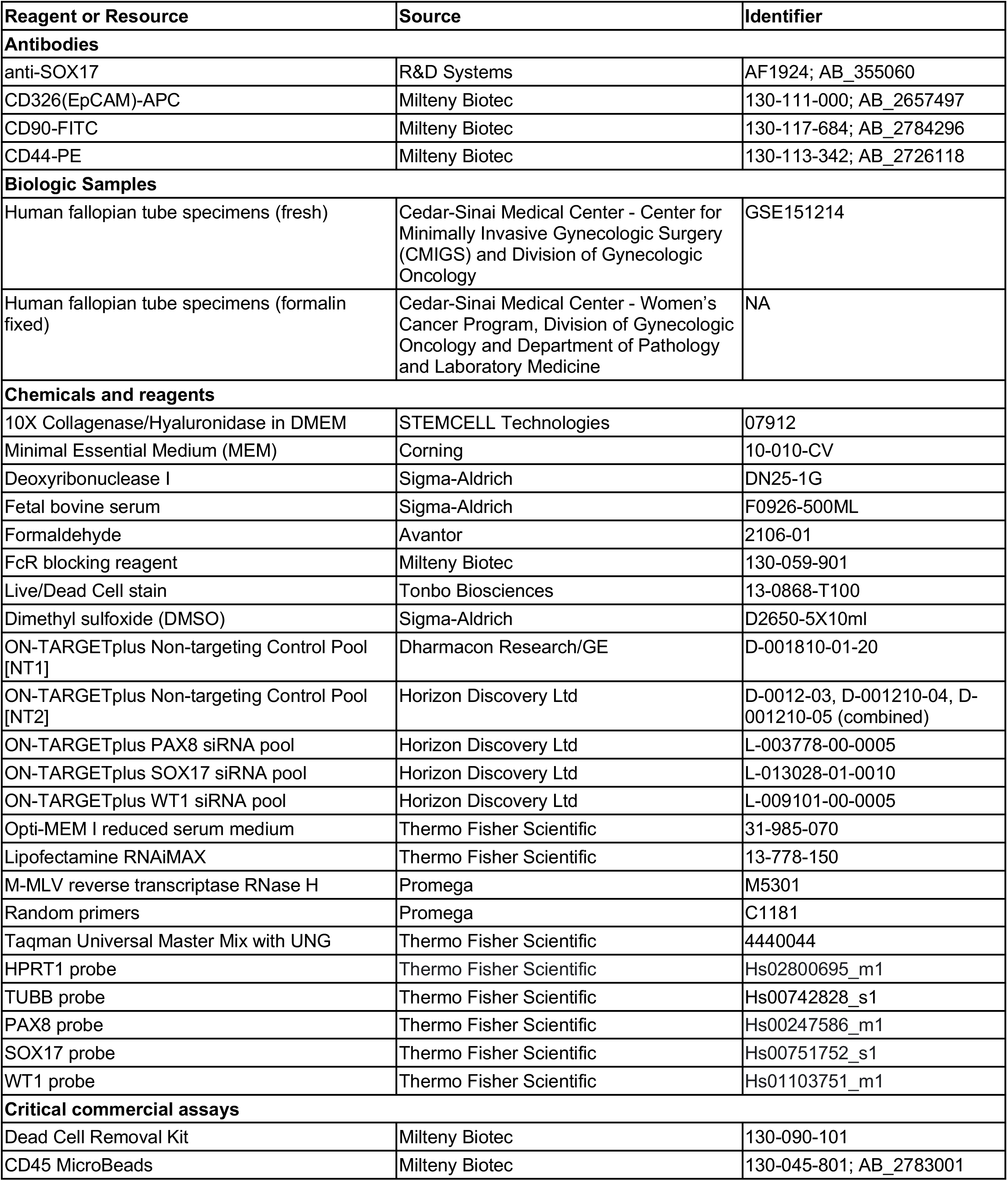
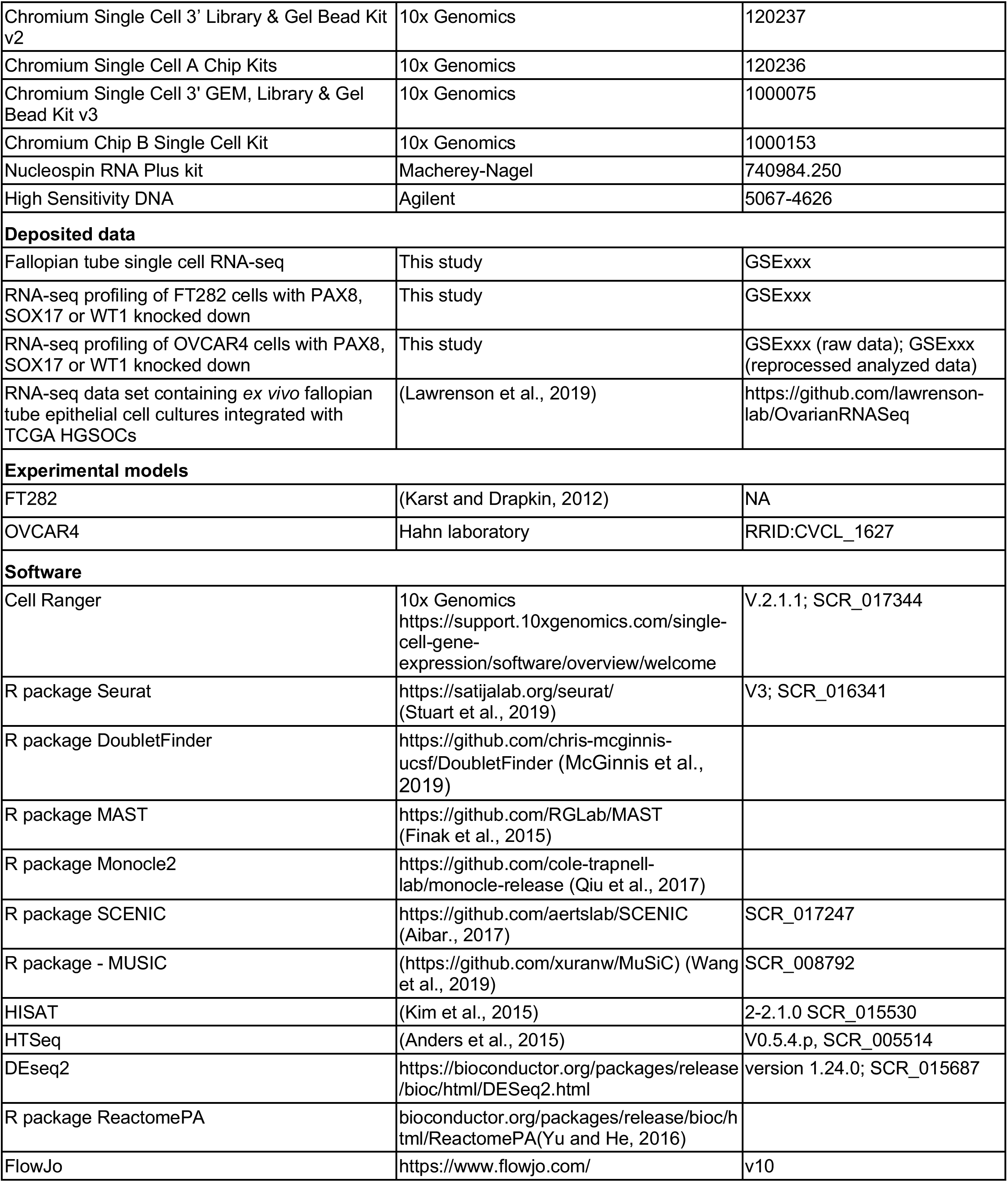

### IRB approval and specimen procurement

This project was performed with approval of the Institutional Review Board at Cedars-Sinai Medical Center (CSMC). All patients provided informed consent. Human fallopian tube tissues were placed in sterile serum-free MEM at 4°C and transferred to the tissue culture laboratory. Pathology review was performed to confirm the absence of malignancy.

### Tissue processing

Primary fallopian tube tissues were transferred into a 10 cm cell-culture dish and the media carefully aspirated. Tissues were minced into ~1-2 mm pieces and digested with 1× Collagenase/Hyaluronidase (STEMCELL Technologies) and 100 μg/mL DNase I (Sigma-Aldrich) in 6.33 mL serum-free MEM. The sample was incubated at 37°C with constant rotation for 90 mins. The supernatant was then harvested, and the remaining tissue flushed with serum free media to collect any remaining cells. The cell suspension was spun at 300 *g* for 10 mins at 4°C. To lyse red blood cells (RBCs), the resulting pellet was resuspended in an RBC lysis buffer (0.8% NH_4_Cl, 0.1% KHCO_3_, pH=7.2) and incubated for 10 mins at room temperature. Cell suspensions were spun again at 300 *g* for 10 mins at 4°C and the cell pellet was resuspended in phosphate buffered saline (PBS), or, if >5% dead cells were observed by trypan blue staining, cells were resuspended in dead cell removal buffer and dead cell removal was performed according to manufacturer’s instructions.

Remaining cells were frozen in 90% fetal bovine serum with 10% DMSO in a Mr. Frosty container placed at - 80°C. Frozen cell vials were stored in LN_2_. Cells were thawed and transferred into a conical tube with 7 mL serum free media and spun at 300 *g* for 10 mins at 4°C. The cell pellet was resuspended in 100 μL PBS. Cells were counted using a hemocytometer and the sample volume adjusted to achieve a cell concentration between 100/μL to 2,000/μL. Due to the low numbers of cells available for the repeat analyses, lower numbers of cells were targeted for capture. For both patients, 10,000 cells were targeted in the fresh capture, but 5,000 cells were targeted in the thawed specimen for patient 6 and 2,000 were targeted in the thawed specimen for patient 7.

### Single cell GEM and barcoding, scRNA-seq library preparation and next-generation sequencing

Single cells are captured and barcoded using 10X Chromium platform (10X Genomics). scRNA-seq libraries were prepared following the instructions from Chromium Single Cell 3ʹ Reagent Kits User Guide (v2 or v3). Briefly, Gel Bead-In EMulsions (GEMs) are generated using single cell preparations. After GEM-RT and cleanup, the cDNA from barcoded single cell RNAs were amplified before quantification using Agilent Bioanalyzer High Sensitivity DNA chips. The single cell 3’ gene expression libraries were constructed and cDNA corresponding to an insertion size of around 350 bp selected. Libraries were quantified using Agilent Bioanalyzer High Sensitivity DNA chip and pooled together to get similar numbers of reads from each single cell before sequencing on a NovaSeq S4 lane (Novogene). For three samples (patients 1-3), capture and NGS were performed by the Cedars-Sinai Applied Genomics, Computation and Translational Genomics Core; for the remaining samples capture and library preparation were performed in-house.

### Single cell RNA-Seq data analysis

#### Preprocessing and integration

Raw reads were aligned to the hg38 reference genome, UMI (unique molecular identifier) counting was performed using Cell Ranger v.2.1.1 (10x Genomics) pipeline with default parameters. We use DoubletFinder (McGinnis et al., 2019) to remove potential doublets based on the expression proximity of each cell to artificial doublets. Only cells with low mitochondrial content (<=10%) were retained for further analysis. To integrate 12 FT scRNA-Seq samples accounting for potential technical variants, we used the anchoring integration method Seurat v3 (Butler et al., 2018; Stuart et al., 2019) which based on canonical correlation (CC) analysis. First, we used *FindIntegrationAnchors* function to find alignment anchors using default 2,000 high variable genes and 30 CC dimensions for defining neighbor search space. Then the anchors were determined and scored for all of 66 possible sample pairs for the integration process using the function *IntegrateData* to account for technical and individual sample variations. Cell type prediction of frozen and cell line single cell data using fresh data as the single cell reference was done using *TransferData* function in Seurat integration pipeline based on defining anchors between two datasets.

### Identification of major cell types and epithelial subgroups

First, we used a standard deviation saturation plot to determine the optimal number of principal components from the integrated dataset. Then, we applied the Louvain community detection-based method which was performed on integrated expression values on shared-nearest-neighbor (SNN) graph as implemented in the *FindCluster* function from the Seurat package for clustering single cell data. Robustness and further clustering analysis were evaluated at the different resolution from 0.1 to 3 with different representative markers to identify the optimal cluster number so that a cluster was called are present at least 2% in at least 3 samples and two close clusters on the cluster tree are merged if there are no differentially expressed gene were detected at cutoff avg_logFC of 0.25 and at least 25% of cells expressed the markers (Differentially expression was performed using MAST (Finak et al., 2015), as implemented in the *FindAllMarker* and *FindMarker* functions in Seurat). The following marker genes were used for global annotation of cell types: Epithelial: *EPCAM*, Keratin genes (*KRT7*, *KRT8*, *KRT18, PAX8, FOXJ1, OVGP1*); T cells: *CD3E*, *CD3D*, *TRBC1/2*, *TRAC*; Myeloid cells: *LYZ*, *CD86*, *CD68*, *FCGR3A*; B cells and plasma cells: *CD79A/B*, *JCHAIN*, *IGKC*, *IGHG3*; Endothelial cells: *CLDN5*; Fibroblasts: *DCN*, *C1R*, *COL1A1*, *ACTA2*; Mast cells: *TPSAB1*. UMAP (uniform manifold approximation and projection) (Becht et al., 2018) was used for visualizing cell-types and clusters with representative markers.

### Identification of markers of epithelial subgroups

Differential expression tests were performed using MAST (Finak et al., 2015), one of the top differential expression analysis methods according to a benchmark study (Soneson and Robinson, 2018) were used to identify markers for each of 10 identified epithelial subsets with average logFC 0.25, expressed in at least 25% of cells in a subset. Pathway analyses were performed using Reactome (Yu and He, 2016).

### Regulatory network inference using SCENIC

We used Single-Cell Regulatory Network Inference And Clustering method, SCENIC (Aibar et al., 2017) to infer regulatory networks, namely regulons, from co-expression of transcription factor and candidate target genes (from RcisTarget database, https://aertslab.org/#scenic). We ran SCENIC for two scenarios: all major cell-types, and within 10 epithelial subsets (the input matrix was normalized expression matrix from Seurat, filtering genes that expressed in at least 10% of samples with minimal count per gene as 3% of the cells, as default in SCENIC). AUC scores for each regulon were used to compare a cluster with the rest to identify the specific regulon and associated transcription factor using a generalized linear model with FDR-corrected p-values < 0.05.

### Pseudotime trajectory analysis using monocle2

We used monocle2 with the DDR tree (Qiu et al., 2017) to infer the potential trajectory of epithelial cells. Epithelial cells were selected to perform pseudo-time ordering based on top 2,000 variable genes after integration (from the *VariableFeature* function in Seurat). Pseudo-time and cell states were inferred using *orderCells* function in monocle2, overlaying the cluster identification from Seurat for consistency. Cell trajectory was plotted using *plot_cell_trajectory* in monocle2 package. Then, we inferred genes associated with the pseudo-time (from early to late) using generalized linear models implemented in function *differentialGeneTest* with parameter “*~sm.ns(Pseudotime)*” using all the genes and RNA expression value (before integration). Hierarchical clustering was then used to infer 4 subset of genes associated with each stage of cellular pseudotime lineage.

### Deconvolution of HGSCs using epithelial subsets

To estimate the potential contribution of epithelial subsets inferred by scRNA-Seq in the bulk sample profiled by RNA-Seq, we used computational deconvolution. First, we defined the subset-specific gene signatures for each of the subset from their differentially expressed genes compared to the rest with the stringent cutoff avg_logFC 1 and adjusted p-value cutoff 0.05. The subsets which shared similar gene signatures after this step were merged leading to the combination of epithelial subsets 2 and 3; as well as the ciliated subsets 1,2,3. Then we used the recent method MuSiC (Wang et al., 2019), accounting for cross-subject gene weighting, to estimate the composition of 6 subsets for normal fallopian tube epithelial *ex vivo* cultures (Lawrenson et al., 2019) and HGSOC samples (Cancer Genome Atlas Research Network, 2011). Enrichment with TCGA molecular subtypes was evaluated by hypergeometric test *phyper* in R.

### Immunohistochemistry

SOX17 immunohistochemistry was performed by the Cedars-Sinai Cancer Biobank and Translational Research Core. Immunohistochemistry was conducted on the Ventan Discovery Ultra Instrument (Roche Ventana). Paraffinized at 72°C with EZ solution (Roche Ventana). Antigen for SOX17 was retrieved via CC1 protocol with prediluted Tris solution, pH 8.0 (Roche Ventana) for 64 minutes at 95°C. This was followed with primary antibody incubation performed with 1:1000 goat anti-SOX17 (R&D) diluted with Antibody Dilution Buffer (Roche Ventana) for 60 minutes at 37°C. Detection was conducted with DISCOVERY UltraMap anti-Goat multimer RUO (Roche Ventana) for 12 minutes at 37°C, and finally with Chromogen DAB CM (Roche Ventana) for 12 minutes at room temperature. For evaluation of SOX17 staining, the following scheme was used: Distribution scoring: 1= 0-24%, 2=25-49%, 3= 50-74%, 4=75-100%; Staining intensity scoring: 0 = absent, 1 = weak, 2 = moderate, 3 = strong. Intensity score reflects the average highest intensity on the slide.

### Validation of progenitor cell populations using flow cytometry

Fallopian tube tissue was obtained from a 48-year-old woman (gravida: 4, para: 2) with a large simple appearing cyst on her left ovary who underwent laparoscopic left salpingo-oophorectomy and right salpingectomy. Bilateral fallopian tubes (fimbria, ampulla, isthmus and infundibulum) were processed into single cells as described above. Dead cell removal was performed and CD45 microbeads used to deplete immune cells. Remaining cells were enumerated and resuspended in PBS containing 2% FBS at a concentration of 4 × 10^7^ cells/ml. Cells were blocked for 15 minutes using an FcR blocking reagent and stained for 30 minutes in a 96-well round bottom plate using EPCAM-APC (1:50 dilution), CD44-PE (1:50 dilution), CD-90-FITC (1:50 dilution) and a live/dead stain. Singly labelled cells were included for compensation, and ‘fluorescence-minus one’ controls included for all stains to enable us to take fluorescence spread into account when placing the gates. Stained cells were washed, fixed with 1.5% formaldehyde, and maintained at 4°C. Acquisition was performed on BD Fortessa cell analyzer cytometer at the Cedars-Sinai Flow Cytometry Core.

### PAX8, SOX17 and WT1 knockdown

OVCAR4 and FT282 cell line models were selected as they most closely recapitulate the cellular and molecular features of human HGSOC (Domcke et al., 2013) and FTSECs, respectively. OVCAR4 cells were cultured in RPMI-1640 supplemented with 10% FBS, 1x NEAA, 11.4 μg/ml of insulin and 1x penicillin/streptomycin. FT282 cells were cultured in DMEM/F12 supplemented with 10% FBS and 1x penicillin/streptomycin. Cultures were maintained at 37°C with 5% CO_2_. Cells were routinely passaged with 0.05% trypsin using standard cell culture procedures. Both cell lines were negative for *Mycoplasma* infections and were authenticated by profiling of short tandem repeats using the Promega Powerplex 16HS assay, performed at the University of Arizona Genomics Core (Table S5).

Cells were reverse transfected using two different non-targeting Control siRNA pools (NT1 and NT2) or pooled oligonucleotides targeting PAX8, SOX17, or WT1 (Horizon Discovery Ltd.) 120 nM of each siRNA pool was diluted in Opti-MEM I (Thermo Fisher Scientific) for 5 minutes, which was then combined with a mix of Opti-MEM I and lipofectamine RNAiMAX (Thermo Fisher Scientific) and incubated for 20 minutes at room temperature. The transfection reagent mix was then combined with 300,000 cells and seeded in a 60mm dish. Media was replenished after 24 hours, and RNA harvested 48 hours later. Cells were washed with cold PBS, collected by scraping and RNA extraction performed using the Nucleospin RNA Plus kit (Macherey-Nagel) according to the manufacturer’s protocol.

### Quantitative PCR

Total RNA was reverse transcribed to cDNA using random primers (Promega) and M-MLV Reverse Transcriptase RNase H (Promega). cDNA was amplified using Taqman Universal Master Mix with UNG (Thermo Fisher Scientific) along with probes for *TUBB, HPRT1* (housekeeping genes) *PAX8*, *SOX17*, or *WT1* (Thermo Fisher Scientific). Data were captured using the QuantStudio 12K Flex Real-Time PCR system (Thermo Fisher Scientific).

### RNA-seq data generation and analysis

Poly-A non-stranded libraries were built from each RNA sample and subjected to 150 bp paired-end sequencing to obtain ~40 million reads/sample. Each knockdown was sequenced in duplicate, using independent knockdown experiments performed at different passages. RNA-seq was performed by BGI using the DNB-seq next generation sequencing platform. Data were filtered and aligned using hisat2-2.1.0 (reference genome: hg38, gencodev26 + ERCC92). A gene-level read count matrix generated using htseq-count (v0.5.4.p3)(Anders et al., 2015). ERCC read counts were removed, and differential gene expression analyses performed using the R package DESeq2 (version 1.24.0). Data for *RUNX3* were extracted (Table S6) and DESeq2 statistics retrieved.

## Data Availability

Data were deposited in NCBI GEO database under the following accession numbers: GSExxx, GSExxx and GSExxx.

## Acknowledgements

This project was supported by an Ovarian Cancer Research Alliance Liz Tilberis Early Career Award (599175) (K.L.), Ovarian Cancer Research Alliance Program Project Development (373356) (B.Y.K., K.L.), a Southern California Clinical and Translational Science Institute Core Voucher (V132) (K.L.) and a Leon Fine Translational Research award (K.L., M.S., K.N.W.). K.L. is also supported by a Research Scholar’s Grant from the American Society (134005). This work was also supported in part by NIH R01 (5R01CA211707, K.L. and S.G.). R.N. is supported in part by a Ruth L. Kirschstein Institutional National Research Service Award (T32) from the NIH (grant number 5 T32 GM 118288-2). The research described was supported in part by NIH/National Center for Advancing Translational Science (NCATS) UCLA CTSI Grant Number UL1TR001881 and in part by Cedars-Sinai Cancer. This research used the following Cedars-Sinai Cores – the Flow Cytometry Core, the Applied Genomics, Computation and Translational (Genomics) Core and the Cancer Biobank and Translational Research Core. We thank the core staff for their support of our research. We thank Dr. Sandra Orsulic and her laboratory for helpful discussions, plus Jenny Lester and study staff from the Women’s Cancer Program biorepository and the Biologic and Epidemiologic Markers of Endometriosis (BEME) Study that provided the specimens used in this study. We also kindly thank the patients that generously donated the specimens used in our research.

## Supplementary Table Legends

**Table S1. Immunohistochemical staining of SOX17 expression in human fallopian tube specimens.** *BRCA1* mutant and wild-type salpingectomy specimens were analyzed. SOX17 epithelial staining distribution and intensity were scored. Staining distribution scoring: 1= 0-24% cells stained, 2=25-49%, 3= 50-74%, 4=75-100%, Staining intensity scoring: 0 = absent, 1 = weak, 2 = moderate, 3 = strong. Intensity score reflects the average highest intensity on the slide. OCP, oral contraceptive pill; HRT, hormone replacement therapy; FT, fallopian tube.

**Table S2. Genes differentially expressed across epithelial clusters.** Genes selected based on average log fold change (logFC) > 0.5, adjusted p-value < 0.01, with a requirement that at least 25% of cells within the cluster expressing the gene of interest.

**Table S3. Enriched pathway analyses in fallopian tube epithelial subsets.** Pathway analyses performed using ReactomePA. There were no pathways enriched in secretory cluster 2. The Gene ID column lists all RefSeq IDs.

**Table S4. Testing for enrichment of tumor clusters within molecular subtypes.** We tested for associations between tumor clusters (defined by unsupervised hierarchical clustering of deconvolution frequencies, with Euclidian distance) and tumor molecular subtypes. Hypergeometric test statistics.

**Table S5. Cell line authentication.** Cell lines were authenticated using the Promega Powerplex 16HS system.

**Table S6.** *RUNX3* **gene expression following knockdown of secretory epithelial cell transcription factors.** Normalized RUNX3 gene expression, in FT282 and OVCAR4, following PAX8, SOX17 and WT1 knockdown. Treatment refers to siRNA target gene. NT, non-targeting control siRNA transfection. Normalisation method is transcripts per million (TPM). Two independent knockdown experiments were performed in each cell line.

## Bibliography

Aibar, S., González-Blas, C.B., Moerman, T., Huynh-Thu, V.A., Imrichova, H., Hulselmans, G., Rambow, F., Marine, J.-C., Geurts, P., Aerts, J., et al. (2017). SCENIC: single-cell regulatory network inference and clustering. Nat. Methods 14, 1083–1086.

Anders, S., Pyl, P.T., and Huber, W. (2015). HTSeq — a Python framework to work with high-throughput sequencing data. Bioinformatics 31, 166–169.

Ardighieri, L., Lonardi, S., Moratto, D., Facchetti, F., Shih, I.-M., Vermi, W., and Kurman, R.J. (2014). Characterization of the immune cell repertoire in the normal fallopian tube. Int. J. Gynecol. Pathol. 33, 581–591.

Barghout, S.H., Zepeda, N., Vincent, K., Azad, A.K., Xu, Z., Yang, C., Steed, H., Postovit, L.-M., and Fu, Y. (2015). RUNX3 contributes to carboplatin resistance in epithelial ovarian cancer cells. Gynecol. Oncol. 138, 647–655.

Becht, E., McInnes, L., Healy, J., Dutertre, C.-A., Kwok, I.W.H., Ng, L.G., Ginhoux, F., and Newell, E.W. (2018). Dimensionality reduction for visualizing single-cell data using UMAP. Nat. Biotechnol.

Butler, A., Hoffman, P., Smibert, P., Papalexi, E., and Satija, R. (2018). Integrating single-cell transcriptomic data across different conditions, technologies, and species. Nat. Biotechnol. 36, 411–420.

Byrd, K.M., Piehl, N.C., Patel, J.H., Huh, W.J., Sequeira, I., Lough, K.J., Wagner, B.L., Marangoni, P., Watt, F.M., Klein, O.D., et al. (2019). Heterogeneity within Stratified Epithelial Stem Cell Populations Maintains the Oral Mucosa in Response to Physiological Stress. Cell Stem Cell 25, 814–829.e6.

Callahan, M.J., Crum, C.P., Medeiros, F., Kindelberger, D.W., Elvin, J.A., Garber, J.E., Feltmate, C.M., Berkowitz, R.S., and Muto, M.G. (2007). Primary fallopian tube malignancies in BRCA-positive women undergoing surgery for ovarian cancer risk reduction. J. Clin. Oncol. 25, 3985–3990.

Cancer Genome Atlas Research Network (2011). Integrated genomic analyses of ovarian carcinoma. Nature 474, 609–615.

Cao, J., Spielmann, M., Qiu, X., Huang, X., Ibrahim, D.M., Hill, A.J., Zhang, F., Mundlos, S., Christiansen, L., Steemers, F.J., et al. (2019). The single-cell transcriptional landscape of mammalian organogenesis. Nature 566, 496–502.

Cheng, L., Lu, W., Kulkarni, B., Pejovic, T., Yan, X., Chiang, J.-H., Hood, L., Odunsi, K., and Lin, B. (2010). Analysis of chemotherapy response programs in ovarian cancers by the next-generation sequencing technologies. Gynecol. Oncol. 117, 159–169.

Chen, F., Gaitskell, K., Garcia, M.J., Albukhari, A., Tsaltas, J., and Ahmed, A.A. (2017). Serous tubal intraepithelial carcinomas associated with high-grade serous ovarian carcinomas: a systematic review. BJOG 124, 872–878.

Crow, J., Amso, N.N., Lewin, J., and Shaw, R.W. (1994). Morphology and ultrastructure of fallopian tube epithelium at different stages of the menstrual cycle and menopause. Hum. Reprod. 9, 2224–2233.

Domcke, S., Sinha, R., Levine, D.A., Sander, C., and Schultz, N. (2013). Evaluating cell lines as tumour models by comparison of genomic profiles. Nat. Commun. 4, 2126.

Donnez, J., Casanas-Roux, F., Caprasse, J., Ferin, J., and Thomas, K. (1985). Cyclic changes in ciliation, cell height, and mitotic activity in human tubal epithelium during reproductive life. Fertil. Steril. 43, 554–559.

Ducie, J., Dao, F., Considine, M., Olvera, N., Shaw, P.A., Kurman, R.J., Shih, I.-M., Soslow, R.A., Cope, L., and Levine, D.A. (2017). Molecular analysis of high-grade serous ovarian carcinoma with and without associated serous tubal intra-epithelial carcinoma. Nat. Commun. 8, 990.

Evans, J., D’Sylva, R., Volpert, M., Jamsai, D., Merriner, D.J., Nie, G., Salamonsen, L.A., and O’Bryan, M.K. (2015). Endometrial CRISP3 is regulated throughout the mouse estrous and human menstrual cycle and facilitates adhesion and proliferation of endometrial epithelial cells. Biol. Reprod. 92, 99.

Finak, G., McDavid, A., Yajima, M., Deng, J., Gersuk, V., Shalek, A.K., Slichter, C.K., Miller, H.W., McElrath, M.J., Prlic, M., et al. (2015). MAST: a flexible statistical framework for assessing transcriptional changes and characterizing heterogeneity in single-cell RNA sequencing data. Genome Biol. 16, 278.

Garcés, J.-J., Simicek, M., Vicari, M., Brozova, L., Burgos, L., Bezdekova, R., Alignani, D., Calasanz, M.-J., Growkova, K., Goicoechea, I., et al. (2020). Transcriptional profiling of circulating tumor cells in multiple myeloma: a new model to understand disease dissemination. Leukemia 34, 589–603.

Ghosh, A., Syed, S.M., and Tanwar, P.S. (2017). In vivo genetic cell lineage tracing reveals that oviductal secretory cells self-renew and give rise to ciliated cells. Development 144, 3031–3041.

Givan, A.L., White, H.D., Stern, J.E., Colby, E., Gosselin, E.J., Guyre, P.M., and Wira, C.R. (1997). Flow cytometric analysis of leukocytes in the human female reproductive tract: comparison of fallopian tube, uterus, cervix, and vagina. Am. J. Reprod. Immunol. 38, 350–359.

Haas, A.R., Tanyi, J.L., O’Hara, M.H., Gladney, W.L., Lacey, S.F., Torigian, D.A., Soulen, M.C., Tian, L., McGarvey, M., Nelson, A.M., et al. (2019). Phase I Study of Lentiviral-Transduced Chimeric Antigen Receptor-Modified T Cells Recognizing Mesothelin in Advanced Solid Cancers. Mol. Ther. 27, 1919–1929.

Healey, A. (2012). Embryology of the female reproductive tract. In Imaging of Gynecological Disorders in Infants and Children, G.S. Mann, J.C. Blair, and A.S. Garden, eds. (Berlin, Heidelberg: Springer Berlin Heidelberg), pp. 21–30.

Hu, Z., Artibani, M., Alsaadi, A., Wietek, N., Morotti, M., Shi, T., Zhong, Z., Santana Gonzalez, L., El-Sahhar, S., KaramiNejadRanjbar, M., et al. (2020). The Repertoire of Serous Ovarian Cancer Non-genetic Heterogeneity Revealed by Single-Cell Sequencing of Normal Fallopian Tube Epithelial Cells. Cancer Cell 37, 226–242.e7.

Karst, A.M., and Drapkin, R. (2012). Primary culture and immortalization of human fallopian tube secretory epithelial cells. Nat. Protoc. 7, 1755–1764.

Karst, A.M., Levanon, K., and Drapkin, R. (2011). Modeling high-grade serous ovarian carcinogenesis from the fallopian tube. Proc Natl Acad Sci USA 108, 7547–7552.

Karst, A.M., Jones, P.M., Vena, N., Ligon, A.H., Liu, J.F., Hirsch, M.S., Etemadmoghadam, D., Bowtell, D.D.L., and Drapkin, R. (2014). Cyclin E1 deregulation occurs early in secretory cell transformation to promote formation of fallopian tube-derived high-grade serous ovarian cancers. Cancer Res. 74, 1141–1152.

Kim, D., Langmead, B., and Salzberg, S.L. (2015). HISAT: a fast spliced aligner with low memory requirements. Nat. Methods 12, 357–360.

Köbel, M., Kalloger, S.E., Boyd, N., McKinney, S., Mehl, E., Palmer, C., Leung, S., Bowen, N.J., Ionescu, D.N., Rajput, A., et al. (2008). Ovarian carcinoma subtypes are different diseases: implications for biomarker studies. PLoS Med. 5, e232.

Kozak, E.L., Palit, S., Miranda-Rodríguez, J.R., Janjic, A., Böttcher, A., Lickert, H., Enard, W., Theis, F.J., and López-Schier, H. (2020). Epithelial Planar Bipolarity Emerges from Notch-Mediated Asymmetric Inhibition of Emx2. Curr. Biol. 30, 1142–1151.e6.

Lawrenson, K., Fonseca, M.A.S., Liu, A.Y., Segato Dezem, F., Lee, J.M., Lin, X., Corona, R.I., Abbasi, F., Vavra, K.C., Dinh, H.Q., et al. (2019). A Study of High-Grade Serous Ovarian Cancer Origins Implicates the SOX18 Transcription Factor in Tumor Development. Cell Rep. 29, 3726–3735.e4.

Lee, C.W.L., Chuang, L.S.H., Kimura, S., Lai, S.K., Ong, C.W., Yan, B., Salto-Tellez, M., Choolani, M., and Ito, Y. (2011). RUNX3 functions as an oncogene in ovarian cancer. Gynecol. Oncol. 122, 410–417.

Lee, Y., Miron, A., Drapkin, R., Nucci, M.R., Medeiros, F., Saleemuddin, A., Garber, J., Birch, C., Mou, H., Gordon, R.W., et al. (2007). A candidate precursor to serous carcinoma that originates in the distal fallopian tube. J. Pathol. 211, 26–35.

Levanon, K., Ng, V., Piao, H.Y., Zhang, Y., Chang, M.C., Roh, M.H., Kindelberger, D.W., Hirsch, M.S., Crum, C.P., Marto, J.A., et al. (2010). Primary ex vivo cultures of human fallopian tube epithelium as a model for serous ovarian carcinogenesis. Oncogene 29, 1103–1113.

Mallette, F.A., Calabrese, V., Ilangumaran, S., and Ferbeyre, G. (2010). SOCS1, a novel interaction partner of p53 controlling oncogene-induced senescence. Aging (Albany NY) 2, 445–452.

McGinnis, C.S., Murrow, L.M., and Gartner, Z.J. (2019). DoubletFinder: Doublet Detection in Single-Cell RNA Sequencing Data Using Artificial Nearest Neighbors. Cell Syst. 8, 329–337.e4.

Medeiros, F., Muto, M.G., Lee, Y., Elvin, J.A., Callahan, M.J., Feltmate, C., Garber, J.E., Cramer, D.W., and Crum, C.P. (2006). The tubal fimbria is a preferred site for early adenocarcinoma in women with familial ovarian cancer syndrome. Am. J. Surg. Pathol. 30, 230–236.

Meserve, E.E.K., Brouwer, J., and Crum, C.P. (2017). Serous tubal intraepithelial neoplasia: the concept and its application. Mod. Pathol. 30, 710–721.

Nevadunsky, N.S., Barbieri, J.S., Kwong, J., Merritt, M.A., Welch, W.R., Berkowitz, R.S., and Mok, S.C. (2009). RUNX3 protein is overexpressed in human epithelial ovarian cancer. Gynecol. Oncol. 112, 325–330.

Ozcan, A., Shen, S.S., Hamilton, C., Anjana, K., Coffey, D., Krishnan, B., and Truong, L.D. (2011). PAX 8 expression in non-neoplastic tissues, primary tumors, and metastatic tumors: a comprehensive immunohistochemical study. Mod. Pathol. 24, 751–764.

Paik, D.Y., Janzen, D.M., Schafenacker, A.M., Velasco, V.S., Shung, M.S., Cheng, D., Huang, J., Witte, O.N., and Memarzadeh, S. (2012). Stem-like epithelial cells are concentrated in the distal end of the fallopian tube: a site for injury and serous cancer initiation. Stem Cells 30, 2487–2497.

Perets, R., Wyant, G.A., Muto, K.W., Bijron, J.G., Poole, B.B., Chin, K.T., Chen, J.Y.H., Ohman, A.W., Stepule, C.D., Kwak, S., et al. (2013). Transformation of the fallopian tube secretory epithelium leads to high-grade serous ovarian cancer in Brca;Tp53;Pten models. Cancer Cell 24, 751–765.

Qiu, X., Mao, Q., Tang, Y., Wang, L., Chawla, R., Pliner, H.A., and Trapnell, C. (2017). Reversed graph embedding resolves complex single-cell trajectories. Nat. Methods 14, 979–982.

Reade, C.J., McVey, R.M., Tone, A.A., Finlayson, S.J., McAlpine, J.N., Fung-Kee-Fung, M., and Ferguson, S.E. (2014). The fallopian tube as the origin of high grade serous ovarian cancer: review of a paradigm shift. J. Obstet. Gynaecol. Can. 36, 133–140.

Reddy, J., Fonseca, M.A.S., Corona, R.I., Nameki, R., Segato Dezem, F., Klein, I.A., Chang, H., Chaves-Moreira, D., Afeyan, L., Malta, T.M., et al. (2019). Predicting master transcription factors from pan-cancer expression data. BioRxiv.

Shimizu, M., Toki, T., Takagi, Y., Konishi, I., and Fujii, S. (2000). Immunohistochemical detection of the Wilms’ tumor gene (WT1) in epithelial ovarian tumors. Int. J. Gynecol. Pathol. 19, 158–163.

Smith, J.M., Wira, C.R., Fanger, M.W., and Shen, L. (2006). Human fallopian tube neutrophils--a distinct phenotype from blood neutrophils. Am. J. Reprod. Immunol. 56, 218–229.

Snegovskikh, V., Mutlu, L., Massasa, E., and Taylor, H.S. (2014). Identification of putative fallopian tube stem cells. Reprod. Sci. 21, 1460–1464.

Soneson, C., and Robinson, M.D. (2018). Bias, robustness and scalability in single-cell differential expression analysis. Nat. Methods 15, 255–261.

Stuart, T., Butler, A., Hoffman, P., Hafemeister, C., Papalexi, E., Mauck, W.M., Hao, Y., Stoeckius, M., Smibert, P., and Satija, R. (2019). Comprehensive Integration of Single-Cell Data. Cell 177, 1888–1902.e21.

Takahashi, K., and Yamanaka, S. (2006). Induction of pluripotent stem cells from mouse embryonic and adult fibroblast cultures by defined factors. Cell 126, 663–676.

Tothill, R.W., Tinker, A.V., George, J., Brown, R., Fox, S.B., Lade, S., Johnson, D.S., Trivett, M.K., Etemadmoghadam, D., Locandro, B., et al. (2008). Novel molecular subtypes of serous and endometrioid ovarian cancer linked to clinical outcome. Clin. Cancer Res. 14, 5198–5208.

Tumbar, T., Guasch, G., Greco, V., Blanpain, C., Lowry, W.E., Rendl, M., and Fuchs, E. (2004). Defining the epithelial stem cell niche in skin. Science 303, 359–363.

Wang, J., Zhao, Y., Wu, X., Yin, S., Chuai, Y., and Wang, A. (2015a). The utility of human fallopian tube mucosa as a novel source of multipotent stem cells for the treatment of autologous reproductive tract injury. Stem Cell Res. Ther. 6, 98.

Wang, S., Zheng, Y., Li, J., Yu, Y., Zhang, W., Song, M., Liu, Z., Min, Z., Hu, H., Jing, Y., et al. (2020). Single-Cell Transcriptomic Atlas of Primate Ovarian Aging. Cell 180, 585–600.e19.

Wang, X., Park, J., Susztak, K., Zhang, N.R., and Li, M. (2019). Bulk tissue cell type deconvolution with multi-subject single-cell expression reference. Nat. Commun. 10, 380.

Wang, Y., Li, L., Wang, Y., Tang, S.N., and Zheng, W. (2015b). Fallopian tube secretory cell expansion: a sensitive biomarker for ovarian serous carcinogenesis. Am. J. Transl. Res. 7, 2082–2090.

White, H.D., Crassi, K.M., Givan, A.L., Stern, J.E., Gonzalez, J.L., Memoli, V.A., Green, W.R., and Wira, C.R. (1997). CD3+ CD8+ CTL activity within the human female reproductive tract: influence of stage of the menstrual cycle and menopause. J. Immunol. 158, 3017–3027.

You, J.-L., Wang, W., Tang, M.-Y., Ye, Y.-H., Liu, A.-X., and Zhu, Y.-M. (2018). A potential role of galectin-1 in promoting mouse trophoblast stem cell differentiation. Mol. Cell. Endocrinol. 470, 228–239.

Yu, G., and He, Q.-Y. (2016). ReactomePA: an R/Bioconductor package for reactome pathway analysis and visualization. Mol. Biosyst. 12, 477–479.

